# UQ-PhysiCell: An extensible Python framework for uncertainty quantification and model analysis in PhysiCell

**DOI:** 10.64898/2026.04.06.716692

**Authors:** Heber L. Rocha, Elmar Bucher, Shuming Zhang, Atul Deshpande, Daniel R. Bergman, Randy Heiland, Paul Macklin

## Abstract

Agent-based models (ABMs) are widely used to study complex multiscale biological systems, particularly in cancer research. However, their high-dimensional parameter spaces, stochasticity, and computational costs pose significant challenges for uncertainty quantification, calibration, and systematic comparison of competing mechanistic hypotheses. PhysiCell has evolved into a growing ecosystem of open-source tools supporting physics-based multicellular modeling, including model construction, visualization, experimental data integration (e.g., spatial multiomics), and downstream output analysis. However, despite these advances, systematic support for uncertainty-aware model analysis, scalable parameter exploration, and formal calibration workflows remains limited. Here, we introduce UQ-PhysiCell, an open-source Python package that enables uncertainty quantification, calibration, and model selection for PhysiCell models using a modular and scalable workflow. UQ-PhysiCell acts as a manager of PhysiCell simulation inputs and outputs, including parameters, initial conditions, rules, and MultiCellDS-compliant objects, and provides automated orchestration of large ensembles of simulations. The framework supports multiple levels of parallelism to accelerate the analysis, including the parallel execution of independent simulations, stochastic replicates, and downstream analysis tasks. UQ-PhysiCell integrates directly with established Python libraries for sensitivity analysis, optimization, Bayesian inference, and surrogate modeling, allowing users to construct customized pipelines that match their modeling goals and computational resource requirements. UQ-PhysiCell decouples model execution from statistical analysis and emphasizes extensibility and reproducibility. This lowers the barrier to rigorous uncertainty-aware analysis and supports systematic evaluation of PhysiCell models in biological and biomedical research.

**Author summary:** We developed UQ-PhysiCell to address a key challenge in agent-based modeling: the systematic quantification of uncertainty in complex stochastic simulations. PhysiCell is widely used to model multicellular biological systems, particularly in cancer research; however, practical tools for uncertainty analysis, calibration, and model comparison are often developed in an ad hoc manner. This makes the results difficult to reproduce and limits the ability to rigorously evaluate competing biological hypotheses. UQ-PhysiCell provides a flexible Python framework that manages the inputs and outputs of PhysiCell simulations and enables large-scale computational analysis. We designed the software to be modular, allowing users to build their own analysis pipelines and combine different methodologies for sensitivity analysis, calibration, and model selection. Rather than enforcing a single workflow, UQ-PhysiCell supports customization to match specific scientific questions and computational requirements. To make uncertainty-aware analyses feasible for computationally intensive agent-based models, UQ-PhysiCell implements multiple parallelism strategies, enabling the concurrent execution of simulations, stochastic replicates, and downstream analyses. UQ-PhysiCell promotes reproducibility, scalability, and methodological flexibility. This helps researchers move beyond single best-fit simulations toward more reliable and interpretable computational modeling.

## Introduction

Mechanistic computational models play a central role in advancing our understanding of complex biological systems by explicitly encoding the biological hypotheses and physical principles. Among these approaches, agent-based models (ABMs) have proven particularly powerful for studying multiscale phenomena in cancer, where individual cell behaviors, microenvironmental interactions, and emergent tissue-level dynamics are tightly coupled [1–7]. As ABMs increasingly inform experimental design and therapeutic strategies, it is essential to rigorously quantify the uncertainty in model predictions and systematically evaluate competing mechanistic assumptions. Recent advancements in spatial multiomics have enabled the development of tools like BIWT [8], which incorporate high-dimensional experimental datasets directly into agent-based simulations. While such approaches strengthen biological grounding, they also increase model complexity and dimensionality, further motivating the need for principled uncertainty quantification and structured model analysis.

Uncertainty quantification (UQ) is especially critical for ABMs due to their large parameter spaces, intrinsic stochasticity, and sensitivity to initial conditions and rule-based interactions. In cancer modeling [9], uncertainties in cellular kinetics, signaling thresholds, microenvironmental properties, and treatment effects can substantially influence simulation outcomes. Without formal UQ, sensitivity analysis, and calibration, model predictions may be difficult to interpret, reproduce, or compare across studies. Despite their importance, these analyses are frequently performed in an ad hoc manner, relying on custom scripts and fragmented tools that limit scalability and reproducibility.

PhysiCell has emerged as one of the leading open-source platforms for physics-based multicellular simulations and is widely adopted by the computational biology and oncology communities [7, 10]. It provides a robust and extensible framework for modeling cell mechanics, diffusion of biochemical substrates, and rule-based cellular behaviors. However, despite its extensibility, it does not natively provide tools for uncertainty-aware analysis, large-scale parameter exploration, or systematic comparison of alternative model formulations. As a result, users must build and maintain their own workflows—managing simulation inputs and outputs, orchestrating high-throughput runs, and performing downstream statistical analyses. This creates barriers to the widespread adoption of sound UQ practices.

To address these challenges, there is a need for a unified and extensible framework that connects PhysiCell simulations with modern uncertainty quantification and model analysis methodologies. Such a framework should simplify model exploration by providing structured management of parameters, initial conditions, and rule sets, while enabling automated generation of parameter sweeps and ensembles of stochastic replicates. Equally important, it should support scalable execution on high-performance computing resources and connect with established analysis tools in the Python ecosystem. UQ-PhysiCell was developed to meet these needs by acting as a Python-based interface and workflow manager for PhysiCell models. The framework organizes simulation inputs and outputs in a consistent and reproducible manner, allowing users to focus on scientific questions rather than low-level file management. The framework couples high-throughput simulation orchestration with flexible analysis pipelines to enable systematic exploration of model behavior across large parameter spaces.

In this work, we introduce UQ-PhysiCell as a modular workflow engine and input/output manager designed specifically for uncertainty-aware analysis of PhysiCell simulations. The framework provides built-in support for sensitivity analysis, calibration, robustness assessment, and model selection, while remaining agnostic to specific statistical methodologies. This design allows users to construct customized pipelines and incorporate alternative analysis techniques as needed. We demonstrate the utility of UQ-PhysiCell through representative examples that highlight how structured workflows and scalable computation improve the reliability, interpretability, and reproducibility of PhysiCell models. UQ-PhysiCell lowers the barrier to rigorous uncertainty quantification. This supports systematic evaluation of mechanistic hypotheses and advances the use of agent-based models as predictive tools in biological and biomedical research.

## Design and Implementation

### Design Philosophy

UQ-PhysiCell’s design rests on four principles: simplicity, modularity, extensibility, and reproducibility. Together, they let the framework support a wide range of scientific workflows without imposing a single rigid pipeline.

Simplicity is achieved by operating directly on standard PhysiCell inputs and outputs, without requiring modifications to model source code. UQ-PhysiCell treats PhysiCell models as black-box dynamical systems and interacts with them exclusively through configuration files and simulation outputs. This design allows users to adopt uncertainty quantification methods incrementally.

UQ-PhysiCell is provided as a Python API, enabling programmatic construction of customized uncertainty-aware workflows. It is also complemented by a graphical user interface (GUI) that lowers the learning curve for new users. The GUI exposes common tasks—such as parameter sweeps, sensitivity analysis, and calibration—without requiring scripting, while advanced users can seamlessly transition to the API for greater flexibility.

Modularity underlies the overall architecture. Simulation orchestration, output processing, model analysis, calibration, surrogate modeling, and model selection are implemented as independent components with well-defined interfaces. Each component wraps existing Python libraries rather than reimplementing them. This separation allows users to extend or replace individual stages without modifying the full pipeline. The same black-box design also lets UQ-PhysiCell interoperate directly with other tools in the PhysiCell ecosystem, such as PhysiCell Studio [11] and BIWT [8].

Reproducibility is enforced through structured data organization, explicit tracking of simulation metadata, and support for MultiCellDS-compliant inputs and outputs [12], ensuring traceability across simulations, replicates, and analyses.

### Software Architecture and Workflow

UQ-PhysiCell follows a workflow with four stages: model specification, high-throughput execution, output processing, and uncertainty-aware analysis (Fig 1).

**Fig 1.**
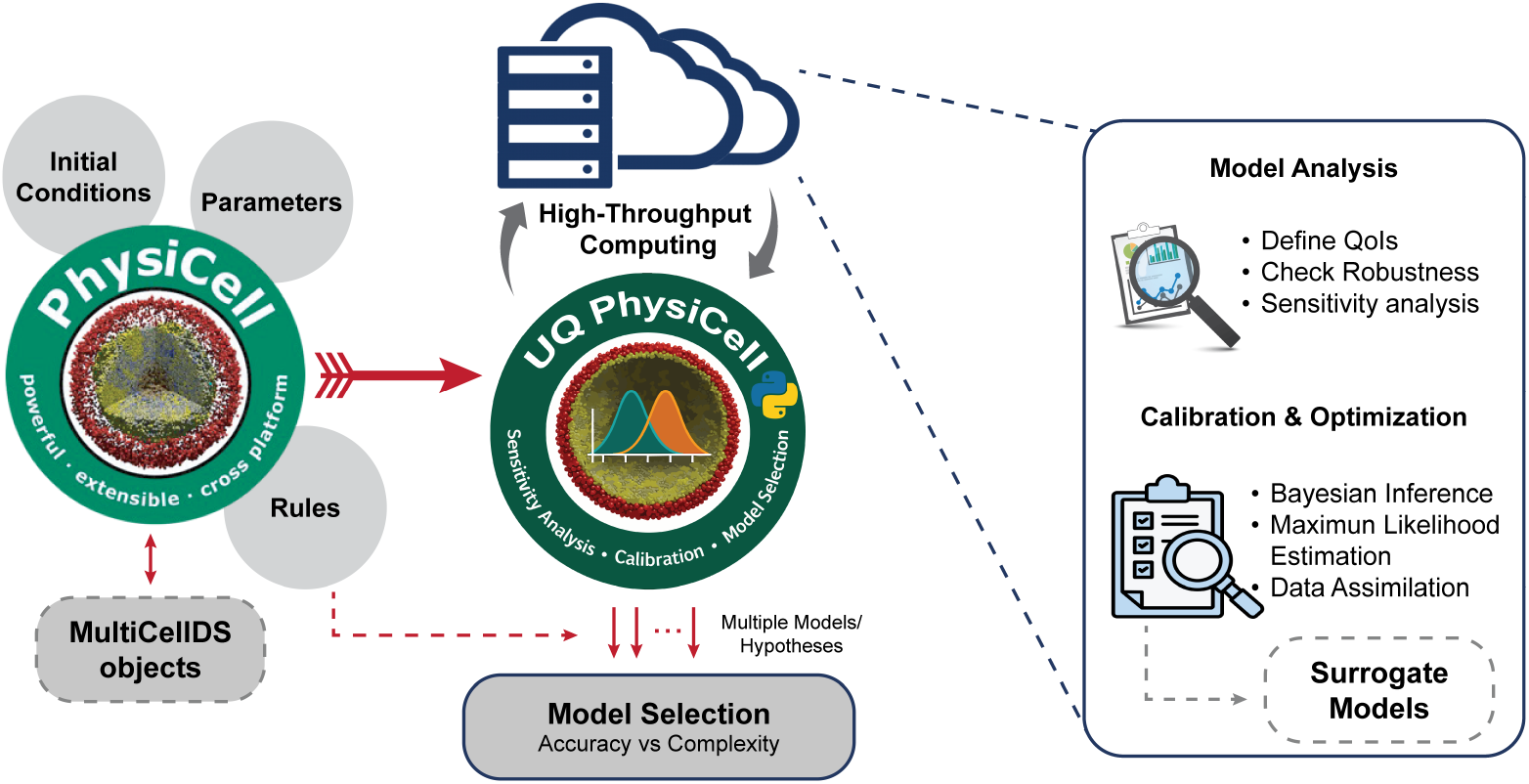
Overview of the UQ-PhysiCell workflow and architecture. UQ-PhysiCell acts as an intermediary between PhysiCell simulations and uncertainty-aware model analysis. Simulation inputs—including parameters, initial conditions, rules, and MultiCellDS objects—are managed and orchestrated through high-throughput computing environments. Outputs from large ensembles of PhysiCell simulations are curated and transformed into analysis-ready data structures, enabling sensitivity analysis, robustness checks, calibration, surrogate modeling, and model selection. The modular design allows users to construct customized pipelines and compare multiple mechanistic hypotheses while balancing predictive accuracy and model complexity.

#### Input specification and model definition

UQ-PhysiCell interfaces directly with PhysiCell model definitions, including initial conditions, parameters, and rule-based behaviors. Users define parameter spaces and experimental conditions using structured Python objects, enabling systematic exploration of high-dimensional parameter spaces.

Support for MultiCellDS objects is provided through integration with the pcdl package [13], allowing standardized representation of multicellular system metadata and experimental context. Each simulation is uniquely defined by a structured specification that includes parameter values, stochastic seeds, and replicate identifiers, ensuring traceability between simulation inputs and downstream results.

#### High-throughput simulation orchestration

UQ-PhysiCell automates the execution of large ensembles of PhysiCell simulations. The framework manages simulation launching, stochastic replicates, and directory organization using consistent and reproducible naming conventions.

To address the computational demands of agent-based modeling, UQ-PhysiCell supports execution across local workstations and HPC clusters. Multiple levels of parallelism enable concurrent execution of parameter samples, replicates, and analysis tasks. Progress tracking and standardized output organization facilitate monitoring of large simulation campaigns and robust recovery from failed runs.

#### Output processing and quantities of interest

PhysiCell simulations generate heterogeneous outputs, including MultiCellDS-compliant XML files, MAT files, and custom logs. UQ-PhysiCell provides an output processing layer that curates and standardizes these raw outputs into analysis-ready data structures, such as Pandas DataFrames.

A central concept in UQ-PhysiCell is the Quantity of Interest (QoI), which defines how complex simulation outputs are reduced for downstream analysis. Formally, a QoI is a mapping

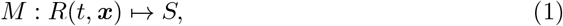

where *R*(*t*, ***x***) represents the full simulation state across time and space, and *S ∈* ℝ*^k^* is a vector of quantities of interest obtained by mapping the raw model output to a reduced set of biologically meaningful summary metrics. QoIs may capture population-level statistics, spatial organization, temporal dynamics, or other biologically relevant features. Standardized QoI extraction enables consistent comparison across parameter sets, stochastic replicates, and competing model formulations.

#### Black-box interaction and stochastic ensembles

UQ-PhysiCell remains agnostic to the internal biological rules implemented within PhysiCell. Accessible parameters are identified by parsing PhysiCell XML configuration files, with each parameter addressed using XPath expressions, allowing users to define a parameter space Θ. For each parameter sample *θ*_*i*_ *∈* Θ, the framework executes an ensemble of *N* stochastic replicates, each initialized with a distinct random seed. This design enables explicit separation of uncertainty arising from parameter variability and intrinsic stochasticity, which is essential for robust uncertainty quantification in agent-based models.

### Core Methodologies and Algorithms

#### Model analysis, robustness, and identifiability

The model analysis layer provides tools for robustness assessment, sensitivity analysis, and model identifiability studies. UQ-PhysiCell supports both global and local sensitivity analysis (SA) to quantify how parameter variations influence selected QoIs.

Beyond sensitivity analysis, UQ-PhysiCell facilitates robustness and identifiability assessments by enabling systematic evaluation of how different QoIs respond to stochastic variability and parameter uncertainty. Users can compare multiple candidate QoIs and quantify their variance across stochastic replicates using metrics such as the relative Monte Carlo standard error (MCSE). This allows identification of QoIs that provide stable and informative summaries of model behavior.

In addition, UQ-PhysiCell enables analysis of variability in the QoI mapping function *M* itself by combining replicate-level variance diagnostics with sensitivity analysis. Users can examine how sensitivity indices change across QoIs and stochastic realizations to assess whether parameters are practically identifiable, and whether observed output variability arises from intrinsic stochasticity or parametric uncertainty.

Global sensitivity analysis is implemented through native integration with the Sensitivity Analysis Library (SALib) [14, 15], supporting variance-based, screening, and distribution-based methods such as Sobol, FAST, Morris, PAWN, and regional sensitivity analysis. UQ-PhysiCell automates sampling scheme generation and routes processed QoIs back to SALib for index computation.

Local sensitivity analysis is performed using a one-factor-at-a-time (OAT) strategy, providing rapid assessment of parameter influence near a nominal parameter set. Custom analysis routines can be implemented through the API to support problem-specific robustness or identifiability studies.

#### Calibration and optimization

UQ-PhysiCell provides a flexible infrastructure for parameter calibration and optimization. Rather than enforcing a single inference paradigm, the framework exposes a modular class-based interface that allows PhysiCell models to be coupled with external optimization and inference libraries.

Built-in support is provided for Bayesian optimization, implemented via BoTorch [16], and Approximate Bayesian Computation (ABC), implemented via pyABC [17]. These methods enable efficient parameter estimation for stochastic models with expensive simulations or intractable likelihoods. The architecture is extensible and designed to support integration with modern simulation-based inference frameworks, including neural posterior estimation approaches [18, 19].

#### Surrogate models and model selection

To mitigate the computational cost of repeated agent-based simulations, UQ-PhysiCell supports the integration of surrogate models within analysis and calibration workflows [20–22]. In its current implementation, surrogate modeling is realized through Gaussian process (GP) models used in Bayesian optimization, enabling sample-efficient exploration of parameter space and accelerated calibration for computationally expensive PhysiCell models.

The software architecture is designed to be surrogate-agnostic, allowing additional surrogate models to be integrated without changes to the core workflow. This includes, but is not limited to, neural network–based surrogates, autoencoder–decoder architectures, and other reduced-order modeling approaches. Such models can be incorporated by extending the existing surrogate interfaces, enabling future acceleration of sensitivity analysis, uncertainty propagation, and inference tasks.

UQ-PhysiCell also provides infrastructure for systematic comparison of competing mechanistic hypotheses, represented by distinct PhysiCell models or alternative rule sets. Model selection is performed by evaluating trade-offs between predictive accuracy and model complexity using information-theoretic or likelihood-based criteria. This design enables principled and reproducible comparison of alternative biological hypotheses while remaining extensible to future surrogate-assisted model selection strategies.

## Results

We demonstrate the capabilities of UQ-PhysiCell through two complementary examples. The first example focuses on a complete end-to-end workflow, illustrating model construction, sensitivity analysis, and parameter calibration for a mechanobiology-driven tumor growth model. The second example highlights a custom analysis use case, where UQ-PhysiCell is used to quantify the sensitivity of individual mechanistic rules in a previously published multicellular invasion model [7]. Together, these examples illustrate how the framework supports both parameter-centric uncertainty quantification and mechanism-centric hypothesis testing in agent-based models.

### End-to-end workflow: model construction, sensitivity analysis, and calibration

We illustrate the proposed UQ-PhysiCell workflow using a simple mechanobiology-driven tumor growth model. Starting from the standard PhysiCell template project, we extend the base model by introducing a mechanofeedback rule on cell proliferation, whereby increased mechanical pressure reduces the probability of cell-cycle entry. In this formulation, the cell-cycle entry rate becomes a spatially and temporally heterogeneous quantity regulated by local mechanical conditions, while the apoptosis rate is kept constant across cells.

Fig 2 summarizes the resulting model behavior. The figure shows representative spatial snapshots of the tumor spheroid over time, colored by the cell-cycle entry rate (panel A) and apoptosis rate (panel B). As the tumor grows and mechanical stresses build up, spatial gradients emerge in cell-cycle entry, with reduced proliferation in the tumor core and higher rates near the periphery, while apoptosis remains spatially uniform by construction.

**Fig 2.**
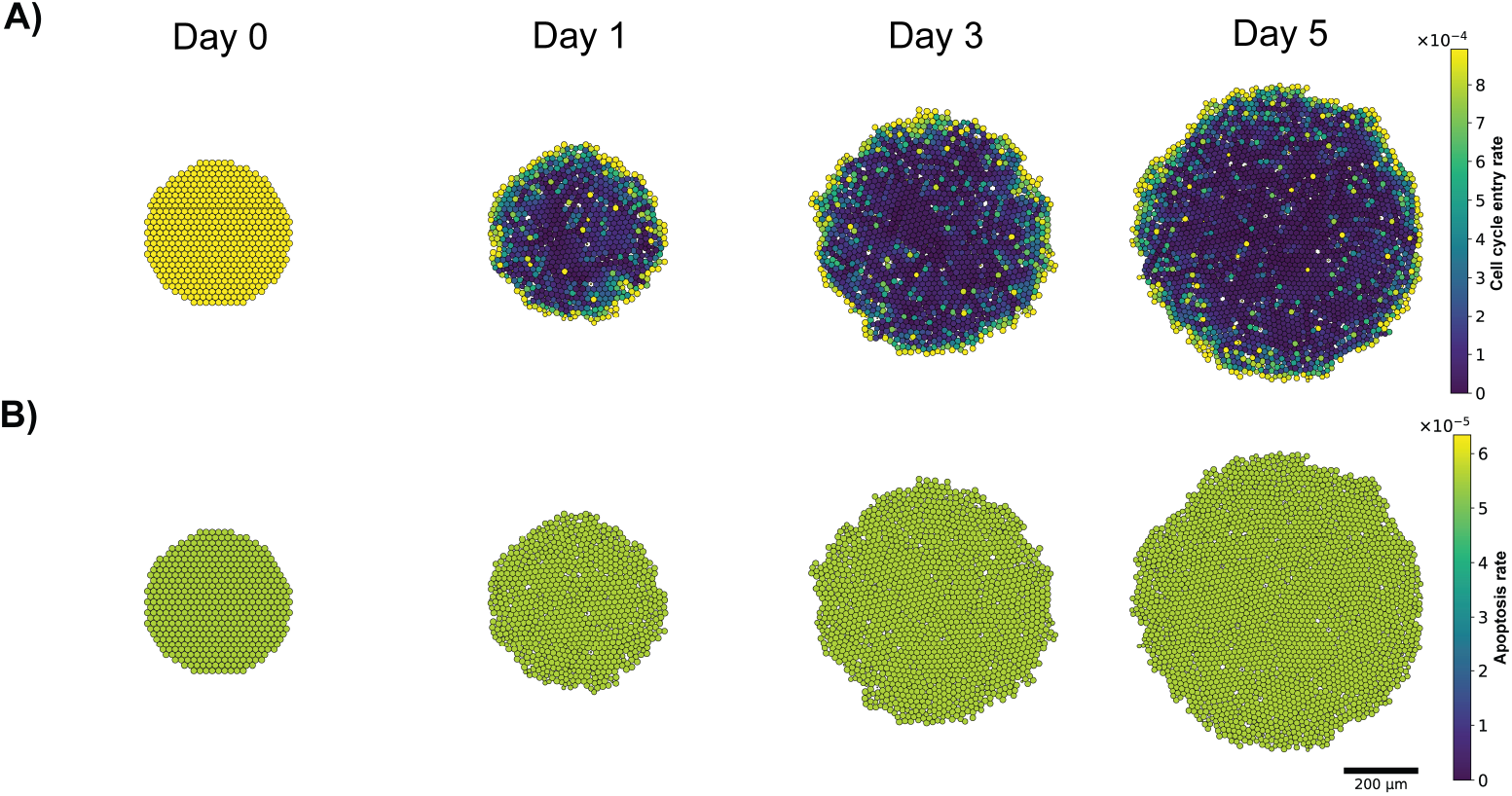
Mechanofeedback-driven tumor growth model behavior. Spatial snapshots of a tumor spheroid simulated with a mechanofeedback-regulated proliferation rule. A: Cell-cycle entry rate at days 0, 1, 3, and 5, showing the emergence of spatial heterogeneity driven by mechanical pressure, with reduced proliferation in the tumor core. B: Apoptosis rate at the same time points, shown here as spatially uniform by construction.

#### Sensitivity analysis and quantity-of-interest selection

After constructing the model, we perform a sensitivity analysis (SA) on two key parameters: cell-cycle duration and apoptosis rate. The analysis is conducted using the UQ-PhysiCell graphical user interface (GUI), enabling systematic exploration of parameter effects without manual scripting. Detailed instructions for performing model construction and sensitivity analysis using the GUI are available in the online documentation (https://uq-physicell.readthedocs.io/en/latest/gui.html).

We evaluate multiple quantities of interest (QoIs), including the live cell population, dead cell population over time, and cumulative cell death at the final time point. Fig 3A shows the ensemble-averaged trajectories and associated relative Monte Carlo standard error (MCSE) for these QoIs across parameter samples. While the dead cell population over time exhibits substantial variability and relatively large relative MCSE, the cumulative cell death displays markedly lower variance across replicates, indicating greater numerical stability and improved identifiability with respect to the selected parameters.

**Fig 3.**
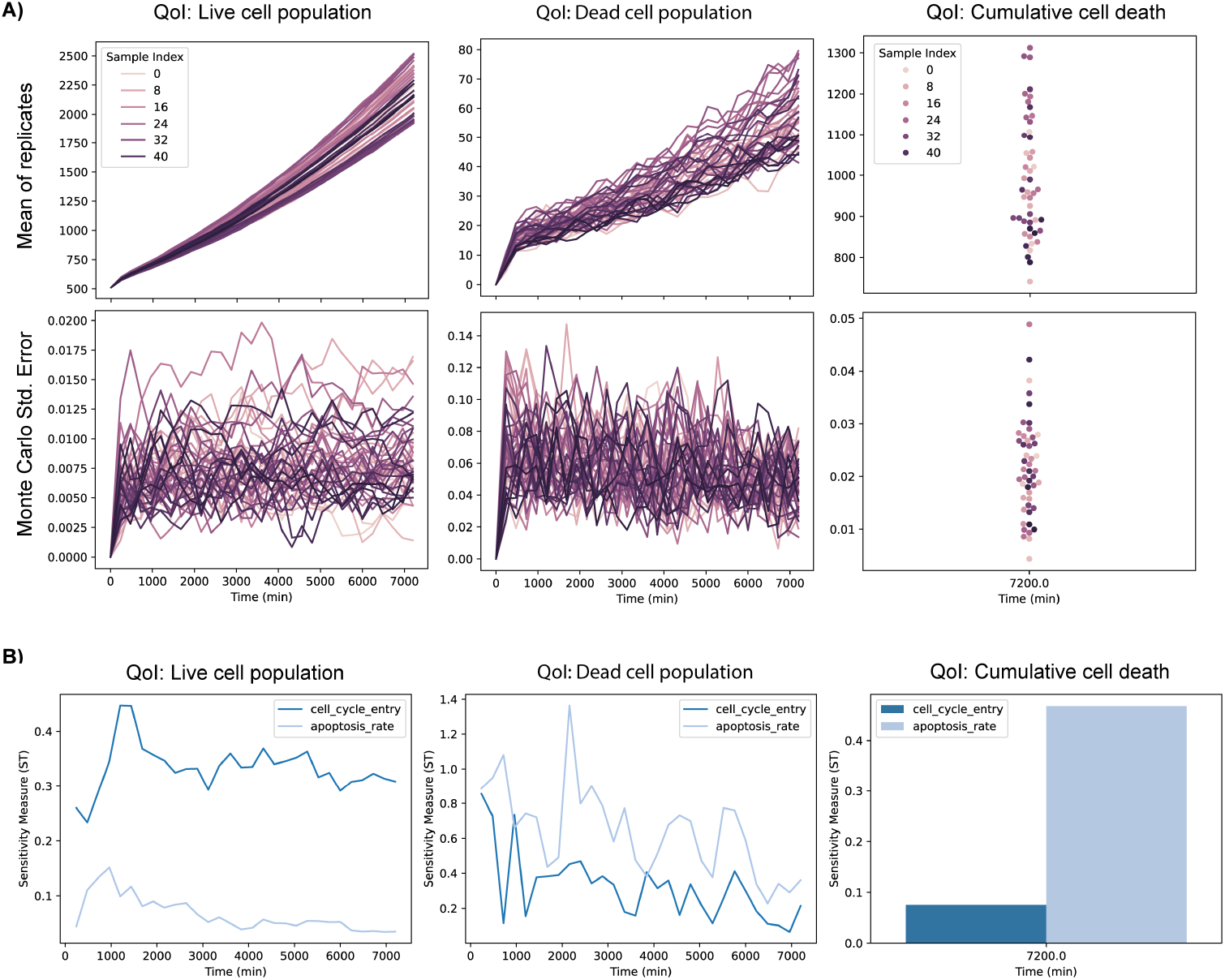
Sensitivity analysis and quantity-of-interest comparison. Sensitivity analysis results for key quantities of interest (QoIs) obtained from ensembles of simulations. A: Ensemble means and relative Monte Carlo standard errors (MCSE) for live cell population, dead cell population over time, and cumulative cell death across parameter samples. Cumulative cell death exhibits substantially lower variance compared to instantaneous dead cell counts. B: Time-resolved total Sobol’ sensitivity index (ST) for cell-cycle duration and apoptosis rate. Cumulative cell death provides clearer parameter discrimination, particularly for apoptosis rate.

This difference is reflected in the sensitivity analysis results shown in Fig 3B. Time-resolved sensitivity indices reveal that cumulative cell death provides a clearer separation of parameter influence—particularly for the apoptosis rate—compared to instantaneous dead cell counts. These results highlight the importance of QoI selection in agent-based model analysis and motivate the use of cumulative cell death as a robust target for subsequent calibration.

#### Bayesian calibration via multi-objective optimization

Following model analysis, we use the simulation database generated during SA to perform model calibration. Synthetic observational data are generated by running the model with a cell-cycle duration of 1 day and an apoptosis rate of 5.787 *×* 10^−5^ *min*^−1^, which serve as ground truth for calibration. Based on the preceding sensitivity analysis, calibration is performed using two quantities of interest (QoIs): (i) the time series of the live cell population and (ii) the cumulative number of cell deaths at the final simulation time.

Bayesian optimization (BO) is initialized using the precomputed simulation database and formulated as a multi-objective optimization problem, targeting simultaneous agreement with the live cell population time series and the cumulative cell death QoIs. Fig 4A shows the resulting Pareto front, consisting of five non-dominated parameter sets that achieve comparable agreement with the target simulation. These Pareto-optimal solutions represent different trade-offs between the calibration objectives (S1 and S2 Figs).

**Fig 4.**
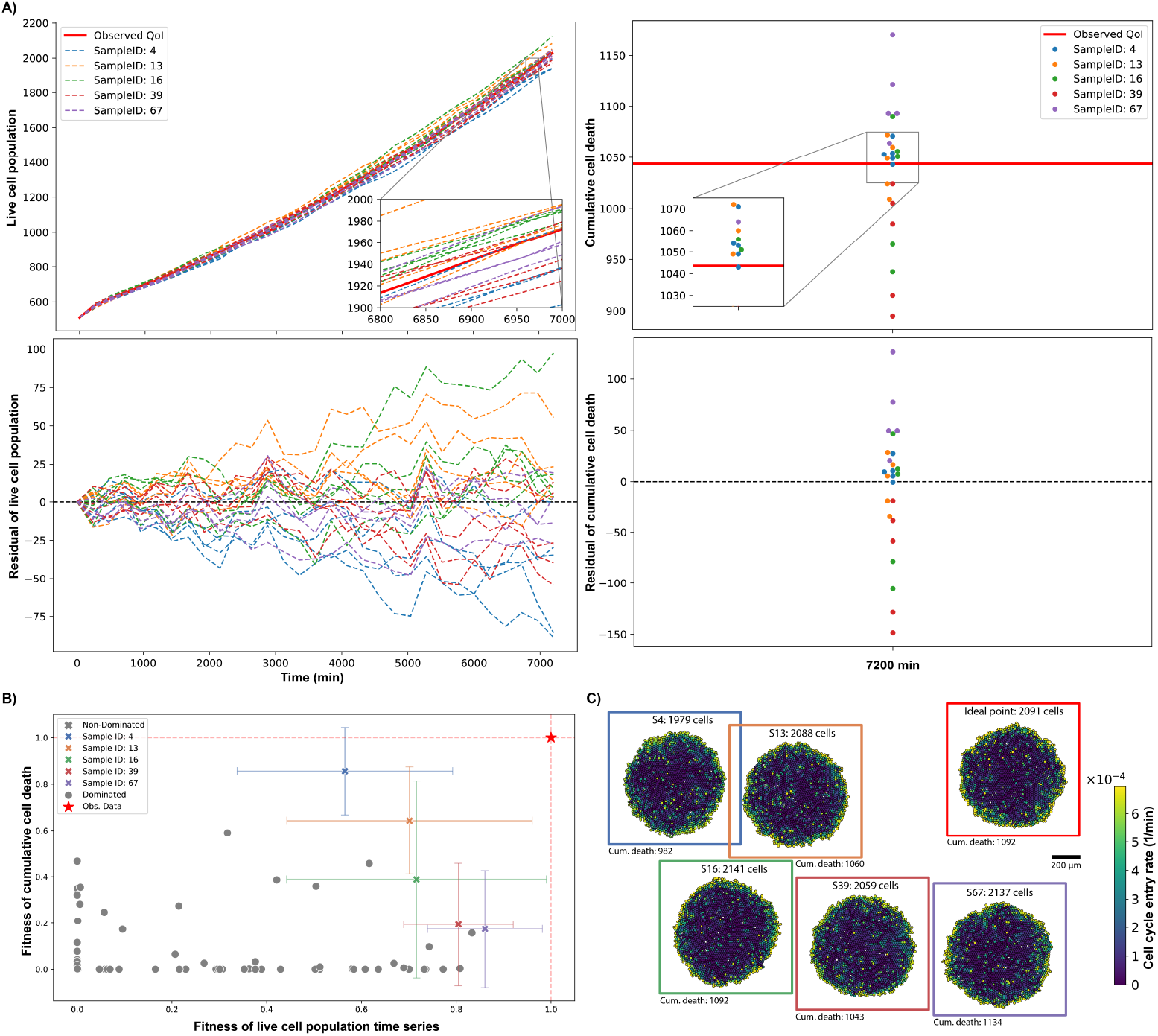
Results of multi-objective Bayesian optimization for model calibration using synthetic data. A: Temporal dynamics of the live cell population (left) and cumulative cell death at the final time point (right) for Pareto-optimal parameter sets, compared against synthetic observational data (red line). The bottom row shows the corresponding residuals for each quantity of interest (QoI). B: Pareto front illustrating the trade-off between fitness objectives. Non-dominated parameter sets (colored ‘x’) represent the optimal balance, error bars indicate the standard deviation of fitness across stochastic replicates, and the red star denotes the theoretical ideal point. C: Spatial snapshots of tumor morphology for selected Pareto-optimal parameter sets. Heatmaps display the cell cycle entry rate, showing consistent macroscopic behavior across top-performing samples.

UQ-PhysiCell provides flexible support for defining custom distance functions, objective weights, and transformations from distance to fitness. Users may specify problem-dependent distance metrics, assign different weights to each QoI, and choose alternative mappings from distance to fitness, enabling tailored calibration strategies across diverse modeling scenarios. In this example, we use the default configuration provided by UQ-PhysiCell. Distances are computed as the sum of squared differences between simulated and observed QoIs. Objective-specific weights are estimated from the empirical ranges of the observational data, so contributions stay balanced across heterogeneous QoIs. The aggregated distance for each objective is then transformed into a fitness value using an exponential mapping,

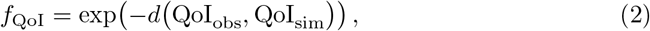

which converts the calibration problem into a maximization task. Under this formulation, the fitness is bounded in [0, 1], with a maximum value of 1 corresponding to perfect agreement between simulation and observation for a given objective.

Fig 4B illustrates how the Pareto-optimal solutions balance the competing fitness objectives, while Fig 4C shows representative final spatial snapshots for each Pareto point in a single replicate. Despite minor differences in spatial organization, all Pareto-optimal solutions reproduce the target QoIs within statistical uncertainty, demonstrating the ability of the workflow to identify multiple plausible parameterizations consistent with the observed data.

### Rule-level sensitivity analysis in a multicellular invasion model

As a second example, we demonstrate the use of UQ-PhysiCell for rule-level sensitivity analysis using a previously developed PhysiCell model of pancreatic cancer invasion mediated by fibroblast–ECM interactions [7]. The model describes interactions between cancer-associated fibroblasts (CAFs), epithelial tumor cells, and mesenchymal tumor cells, and includes multiple mechanistic rules governing cell migration, phenotypic transitions, and contact inhibition.

To assess the contribution of individual rules to model behavior, we performed a one-rule-at-a-time perturbation analysis. Starting from the nominal model in which all rules are active, we systematically inactivated a single rule and quantified deviations in model outputs relative to the full model (S1 Table). Simulations were performed across seven initial conditions (ICs), defined by different epithelial tumor cell–to–CAF seeding ratios, while maintaining a total of 1,000 cells per simulation. Each scenario was simulated for seven days with multiple stochastic replicates.

Fig 5A illustrates the seven initial conditions and representative simulation outcomes at day 7, highlighting distinct invasion patterns and spatial organization driven by cell–cell and cell–ECM interactions. Fig 5B reports the mean behavior across replicates for selected quantities of interest (QoIs), including population dynamics and radial expansion of each cell type, demonstrating a strong dependence of model outputs on the chosen initial condition.

**Fig 5.**
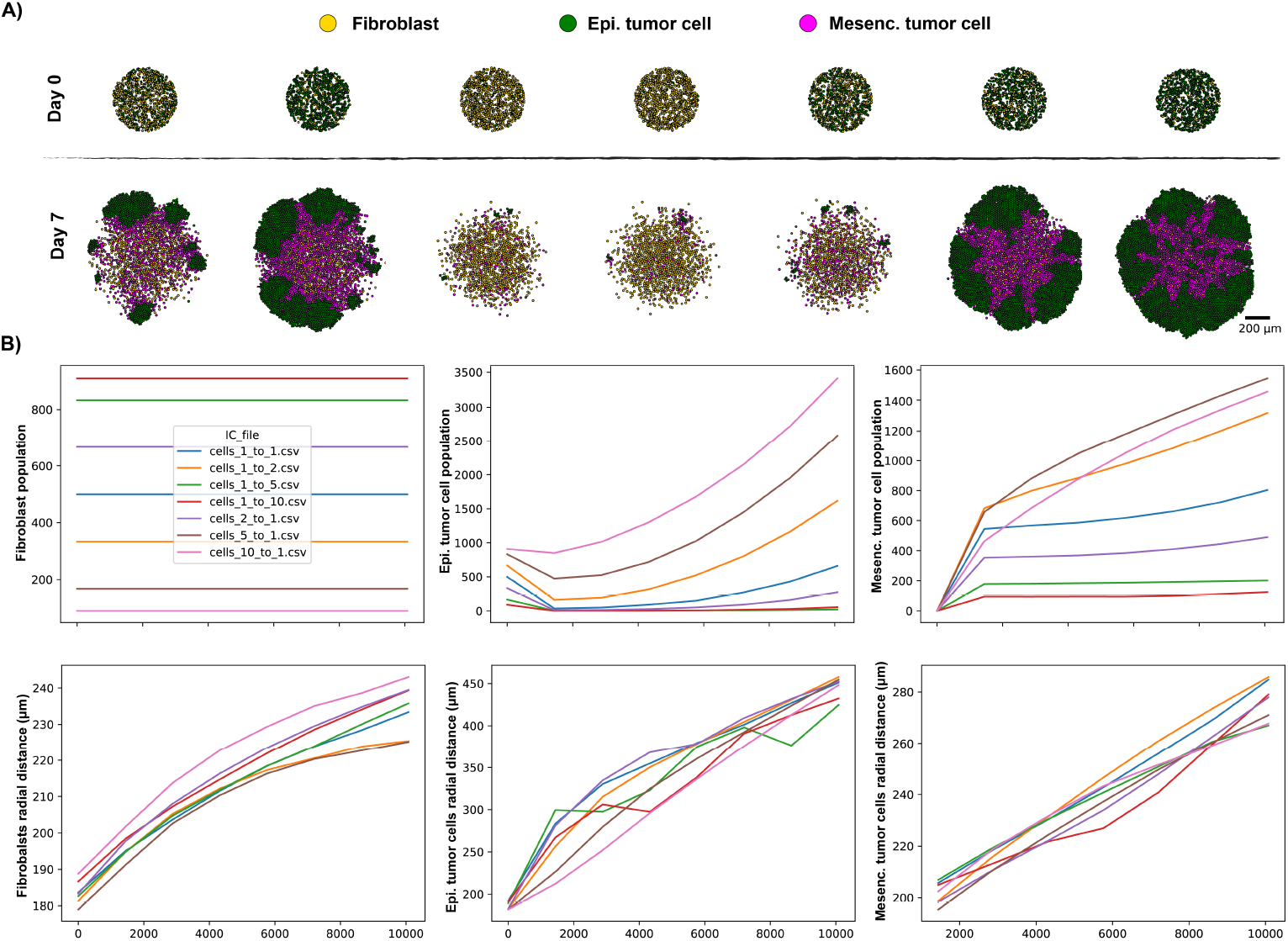
Initial conditions and baseline dynamics of the invasion model. A: Seven initial conditions defined by different epithelial tumor cell–to–CAF seeding ratios, shown at day 0 (top) and representative simulation outcomes at day 7 (bottom). B: Mean behavior across stochastic replicates for selected quantities of interest, including population dynamics and radial expansion of each cell type, illustrating the strong dependence of model outputs on initial condition.

For each IC, we quantified the sensitivity of individual rules by computing the root mean square deviation (RMSD) between the QoIs obtained from rule-inactivated simulations and the nominal model (S3-S7 Figs). These RMSD values provide a direct measure of how strongly each rule influences the selected QoIs under a given IC. To obtain a global measure of rule importance, RMSDs were normalized per QoI and IC, and averaged across features (Fig 6A).

**Fig 6.**
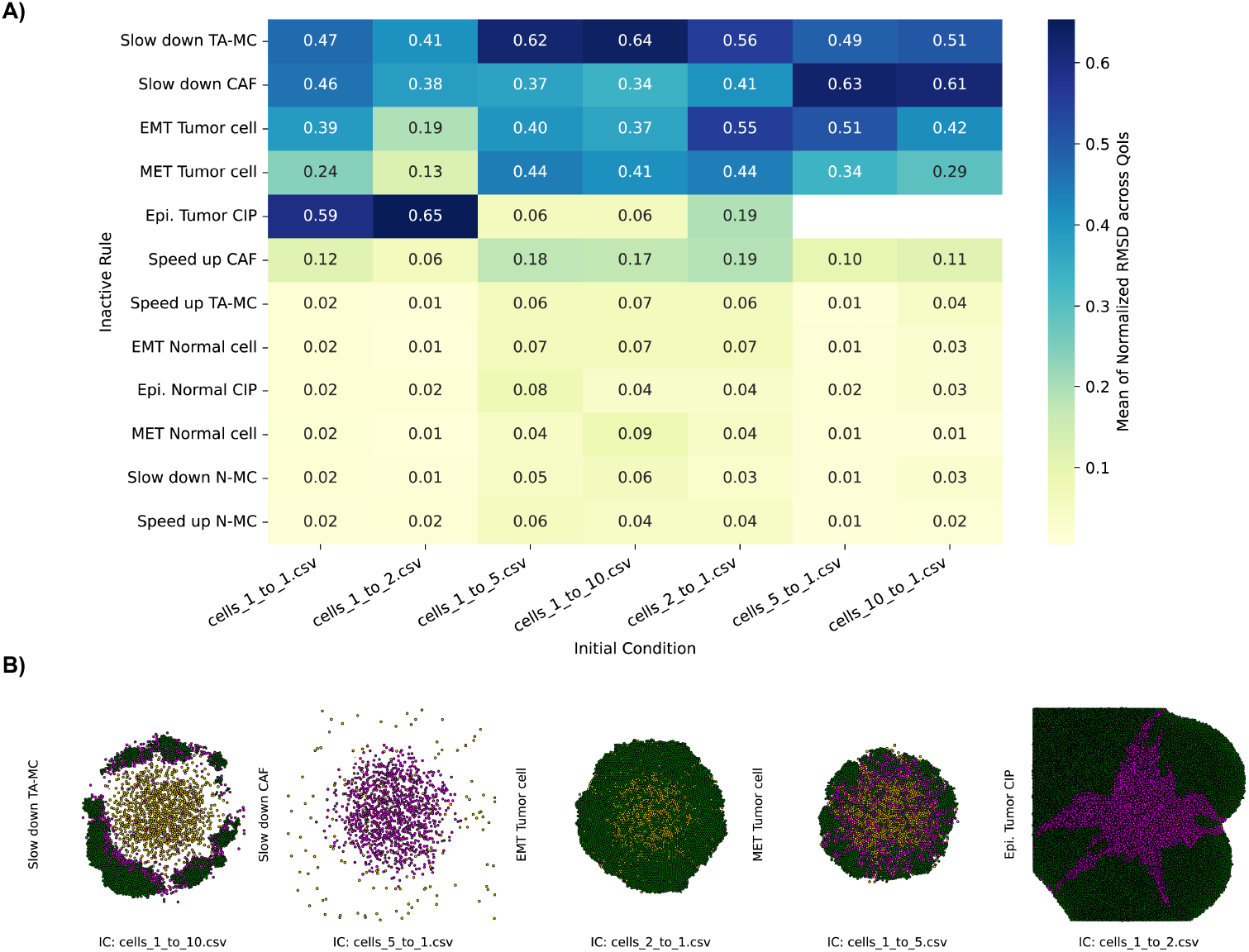
Rule-level sensitivity analysis across initial conditions. A: Global sensitivity of individual rules computed by averaging normalized root mean square deviation (RMSD) across quantities of interest and initial conditions. Higher values indicate stronger deviations from the nominal model when a rule is inactivated. B: Representative spatial snapshots at day 7 for selected high-impact rule perturbations, illustrating how inactivation of specific rules leads to distinct invasion morphologies depending on the initial condition.

This analysis reveals that rule importance is highly context-dependent, varying substantially across initial conditions. Nevertheless, two rules—those associated with slowing down tumor-associated mesenchymal cells (TA-MC) and slowing down cancer-associated fibroblasts (CAFs)—exhibit consistently high influence across all initial conditions. In contrast, rules governing epithelial–mesenchymal transition (EMT), inhibition of mesenchymal–epithelial transition (MET), and contact inhibition of proliferation in epithelial tumor cells show pronounced effects only in specific initial condition scenarios. By comparison, rules that increase motility of CAFs and TA-MCs have little impact on the selected QoIs. The same holds for rules involving normal epithelial and normal mesenchymal cells (N-MC), since these populations aren’t present in the simulated system. Fig 6B illustrates representative spatial outcomes at day 7 for selected high-impact rule perturbations, emphasizing how local rule changes can lead to qualitatively distinct invasion patterns (S8 Fig).

The limited impact observed for rules that increase the motility of cancer-associated fibroblasts and tumor-associated mesenchymal cells may indicate missing mechanisms in the current extracellular matrix (ECM) representation. In particular, ECM adhesion, fiber alignment, and directional guidance cues often strongly influence cell motility in invasive tumors, but the current model does not explicitly capture these mechanisms. Incorporating a more detailed ECM description—such as fiber orientation and anisotropic adhesion—may therefore alter the sensitivity of motility-related rules. Such extensions could be implemented using the PhysiMeSS add-on [23], which enables explicit modeling of ECM structure and cell–matrix interactions.

In contrast, inactivation of the contact inhibition of proliferation rule in epithelial tumor cells leads to unbounded cell proliferation without spatial constraints. This behavior results in overcrowded cell configurations and exponential population growth, as illustrated in Fig 6B for the scenario in which the epithelial tumor contact inhibition rule is disabled under the `cells_1_to_2` initial condition. In more extreme initial conditions (`cells _5 to_1` and `cells_10_to_1`), this unconstrained growth caused simulations to exceed the imposed runtime limit of 1.5 hours before reaching the final simulation time. These outcomes show that rule-level perturbations can expose both biological sensitivities and computational failure modes. This underscores the importance of systematic rule testing in complex agent-based models.

## Availability and Future Directions

Comprehensive documentation, including installation instructions, tutorials, and a collection of worked examples, is available at https://uq-physicell.readthedocs.io. These resources enable users and reviewers to reproduce the results presented in this manuscript and to apply UQ-PhysiCell to new modeling studies. The project, UQ-PhysiCell, is openly available with archived releases and reproducibility resources provided through https://doi.org/10.5281/zenodo.17823176. The software and accompanying documentation are intended to facilitate transparent and reproducible workflows for uncertainty-aware analysis of agent-based models developed with PhysiCell [10].

UQ-PhysiCell is implemented entirely in Python (≥ 3.10) and distributed as free, open-source software under the BSD 3-Clause license. Source code, version history, and issue tracking are hosted at https://github.com/heberlr/UQ_PhysiCell, with tagged releases additionally published to the Python Package Index (PyPI) as uq physicell, enabling installation through standard pip tooling. Core dependencies (NumPy, pandas, SciPy, and SALib for sensitivity analysis) are installed automatically, while optional extras enable Bayesian optimization (BoTorch, GPyTorch) and Approximate Bayesian Computation (pyABC, Dask) as needed. Code correctness is maintained through a continuous integration pipeline (GitHub Actions) that lints the codebase and executes the unit test suite with coverage reporting on every pull request, supporting the reliability of releases distributed through PyPI and Zenodo. There are no restrictions on use by academic or non-academic users.

UQ-PhysiCell provides a structured interface for uncertainty quantification, sensitivity analysis, and calibration. This supports more rigorous and reproducible computational studies. The framework promotes community standards for sharing, benchmarking, and systematically evaluating PhysiCell models, while bridging agent-based modeling workflows with modern statistical inference and data-driven analysis tools. Rather than proposing new statistical methodology or competing with established UQ and inference libraries, UQ-PhysiCell is deliberately designed as an integrative layer. It absorbs the engineering burden of simulation orchestration, output curation, and interface management, decreasing the friction of coupling PhysiCell models with mature, general-purpose analysis ecosystems such as SALib, BoTorch, and pyABC, rather than duplicating their functionality.

Despite these capabilities, several limitations remain. Currently, the simulation database is managed locally, which can restrict collaborative workflows and large-scale distributed studies. In addition, careful selection of quantities of interest (QoIs) remains necessary to ensure robust model analysis; the integration of automated QoI discovery methods is an important area for future work [24]. Finally, surrogate model fidelity may depend on the diversity and coverage of training simulations, particularly when surrogate models are used to accelerate calibration or uncertainty propagation [25].

Moving forward, we aim to expand the UQ-PhysiCell ecosystem to support more complex modeling paradigms and collaborative workflows. A primary objective is the implementation of a federated execution layer that allows simulations to run across multiple heterogeneous clusters while asynchronously aggregating results into a centralized, schema-agnostic NoSQL database. This document-oriented approach will efficiently manage heterogeneous agent-based data—where simulation outputs vary in dimensionality and metadata—without the limitations of rigid relational table structures or file-locking conflicts inherent in local databases.

To improve the interpretability of these large-scale ensembles, we plan to develop integrated visualization dashboards for real-time monitoring of calibration progress and sensitivity indices. Furthermore, we will extend the framework to natively support multiscale extensions such as PhysiBoSS [26], which integrates stochastic Boolean signaling networks with agent-based dynamics. This will streamline the analysis of hybrid models, keeping uncertainty quantification robust even as simulations incorporate complex intracellular logic and disparate spatial scales. Finally, we envision a deeper coupling with PhysiCell Studio [11], creating a seamless bridge between model design, graphical parameter exploration, and rigorous statistical analysis.

Furthermore, the framework can naturally interoperate with emerging tools in the PhysiCell ecosystem. For example, PhysiGym [27] provides an interface between Physi-Cell simulations and reinforcement learning environments through the Gymnasium API, enabling learning-based control strategies to be applied to multicellular simulations. Integration with such tools may enable future workflows where reinforcement learning agents interact with UQ-driven simulation ensembles for adaptive experimental design and model exploration.

Beyond classical machine learning surrogates and reinforcement learning agents, recent advances in large language models (LLMs) suggest complementary directions for the UQ-PhysiCell framework. LLM-assisted calibration could involve proposing informative parameter priors or distance-function weightings from natural-language descriptions of the target biology, and generating human-readable interpretations of sensitivity indices, Pareto fronts, and calibration outcomes to lower the barrier for non-expert users. Similarly, the automated QoI discovery challenge noted above could be approached by coupling LLM-based reasoning over simulation outputs and metadata with representation-learning techniques to propose candidate summary statistics for emergent spatial or temporal patterns not anticipated a priori. These LLM-assisted capabilities remain unexplored in the current implementation and would require careful validation against domain expert judgment before adoption in rigorous UQ workflows, but the framework’s modular, black-box design—already decoupling simulation execution from analysis—is well positioned to incorporate such agentic components as the tooling matures.

## Supporting information

**S1 Fig. Normalized parameter samples and Pareto optimal points**. Visualization of the parameter space explored during Bayesian Optimization (BO) for the mechanobiology-driven tumor growth model. Each point represents a parameter set sampled by the optimizer, with the Pareto points highlighted to show their proximity to the reference values.

**S2 Fig. Objective function landscape and Pareto points**. Scatter plots of parameter values against the fitness scores for each quantity of interest (QoI). This figure illustrates the sensitivity of the fitness functions to specific parameters and identifies the Pareto points that minimize error across multiple model outputs. The dashed red lines indicate the reference values used to generate the observational data.

**S1 Table. Mechanistic rules of the invasion model**. Each rule is defined specifically for a given cell type. Following the standardized PhysiCell grammar, each rule is modeled as a behavioral response to a microenvironmental signal. The response is mathematically defined by a Hill function, where a specific signal (e.g., local pressure or ECM density) increases or decreases a target behavior from a base value toward a saturation value. Each functional relationship is characterized by four key parameters: the base and saturation levels, the half-max, and the Hill power, which dictates the sensitivity or “switch-like” nature of the response.

**S3 Fig. Normalized RMSD for a population of epithelial tumor cells**. The normalized Root Mean Square Deviation (RMSD) between each model output under varying initial conditions and rule inactivations relative to the nominal scenario. This figure quantifies the impact of inactivating each rule on model behavior compared to the number of epithelial tumor cells in the nominal scenario.

**S4 Fig. Normalized RMSD for a population of mesenchymal tumor cells**. The normalized Root Mean Square Deviation (RMSD) between each model output under varying initial conditions and rule inactivations relative to the nominal scenario. This figure quantifies the impact of inactivating each rule on model behavior compared to the number of mesenchymal tumor cells in the nominal scenario.

**S5 Fig. Normalized RMSD for radial distance of fibroblasts**. The normalized Root Mean Square Deviation (RMSD) between each model output under varying initial conditions and rule inactivations relative to the nominal scenario. This figure quantifies the impact of inactivating each rule on model behavior compared to the radial distance of fibroblasts in the nominal scenario.

**S6 Fig. Normalized RMSD for radial distance of epithelial tumor cells**. The normalized Root Mean Square Deviation (RMSD) between each model output under varying initial conditions and rule inactivations relative to the nominal scenario. This figure quantifies the impact of inactivating each rule on model behavior compared to the radial distance of epithelial tumor cells in the nominal scenario.

**S7 Fig. Normalized RMSD for radial distance of mesenchymal tumor cells**. The normalized Root Mean Square Deviation (RMSD) between each model output under varying initial conditions and rule inactivations relative to the nominal scenario. This figure quantifies the impact of inactivating each rule on model behavior compared to the radial distance of mesenchymal tumor cells in the nominal scenario.

**S8 Fig. Spatial snapshots of nominal and high-impact scenarios**. Final simulation snapshots (last time point) across different initial conditions. The first column displays the nominal (baseline) behavior, while columns 2–5 visualize the scenarios with the highest impact on model outputs, ordered by their influence as determined by the sensitivity analysis.

## Acknowledgments

We thank members of the Fertig Lab and the Forecasting Group for valuable discussions and feedback during the development of this software. We also thank the PhysiCell community for ongoing discussions and insights that motivated and informed the design of UQ-PhysiCell. Part of the computational simulations of this research were performed on high-throughput computing clusters at Indiana University, which were supported in part by Lilly Endowment, Inc., through its support for the Indiana University Pervasive Technology Institute.

## Notes

https://uq-physicell.readthedocs.io

https://github.com/heberlr/UQ_PhysiCell

### Competing Interest Statement

The authors have declared no competing interest.

### Summary of Updates

This revision consists of clarity and transparency improvements; no changes were made to the methods, results, figures, or conclusions. - Expanded the Availability and Future Directions section with explicit software engineering details: license (BSD 3-Clause), source repository (GitHub), PyPI installation, core/optional dependencies, and continuous integration/testing infrastructure. - Added a paragraph to Future Directions discussing potential future integration of LLM-assisted calibration/interpretation and automated quantity-of-interest (QoI) discovery, explicitly noted as unexplored directions requiring future validation. - Clarified the Abstract's description of the PhysiCell tool ecosystem, distinguishing model construction/visualization, experimental data integration (e.g., spatial multiomics), and downstream output analysis as separate categories. - Added an explicit statement positioning UQ-PhysiCell as an integrative layer over existing UQ/inference libraries rather than a competing implementation. - General copyediting pass throughout for sentence clarity and consistency.

### Funding

- National Institutes of Health: NIH/NCI U24CA284156
- Jayne Koskinas Ted Giovanis Foundation

