## Supporting information for "UQ-PhysiCell: An extensible Python framework for uncertainty quantification and model analysis in PhysiCell"

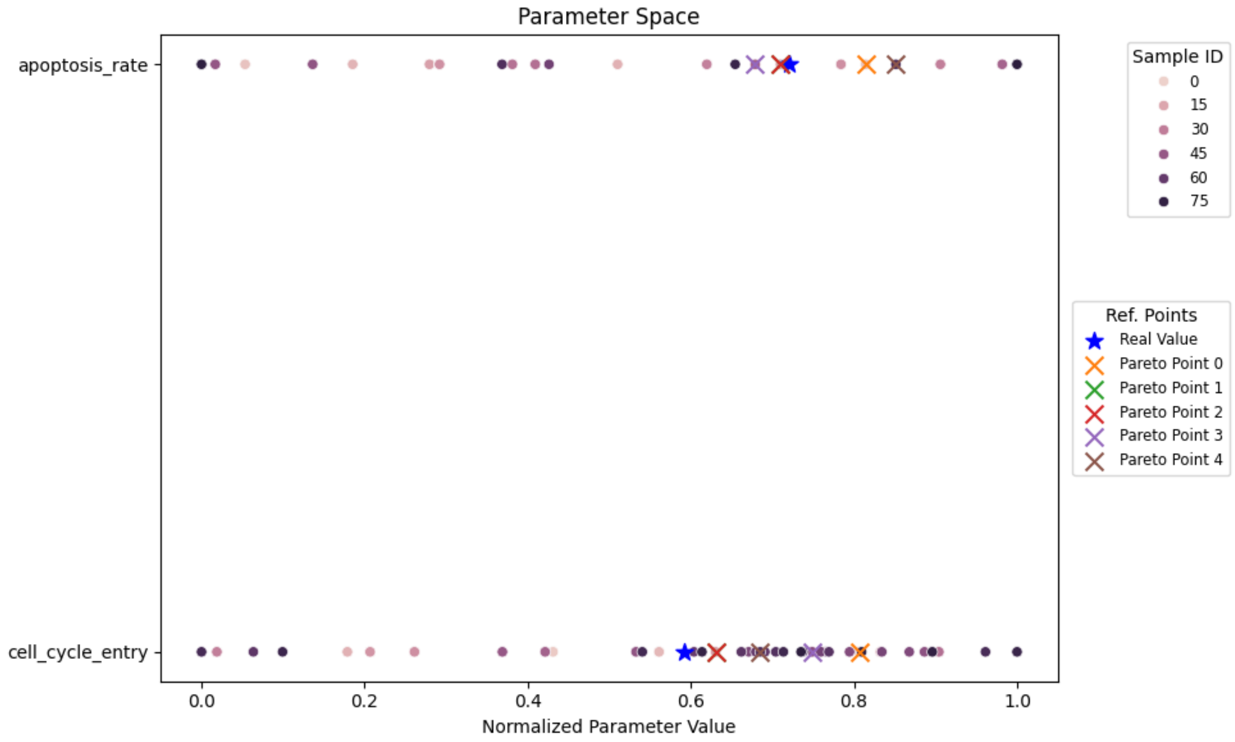


**S1 Fig. Normalized parameter samples and Pareto optimal points**. Visualization of the parameter space explored during Bayesian Optimization (BO) for the mechanobiology-driven tumor growth model. Each point represents a parameter set sampled by the optimizer, with the Pareto points highlighted to show their proximity to the reference values.

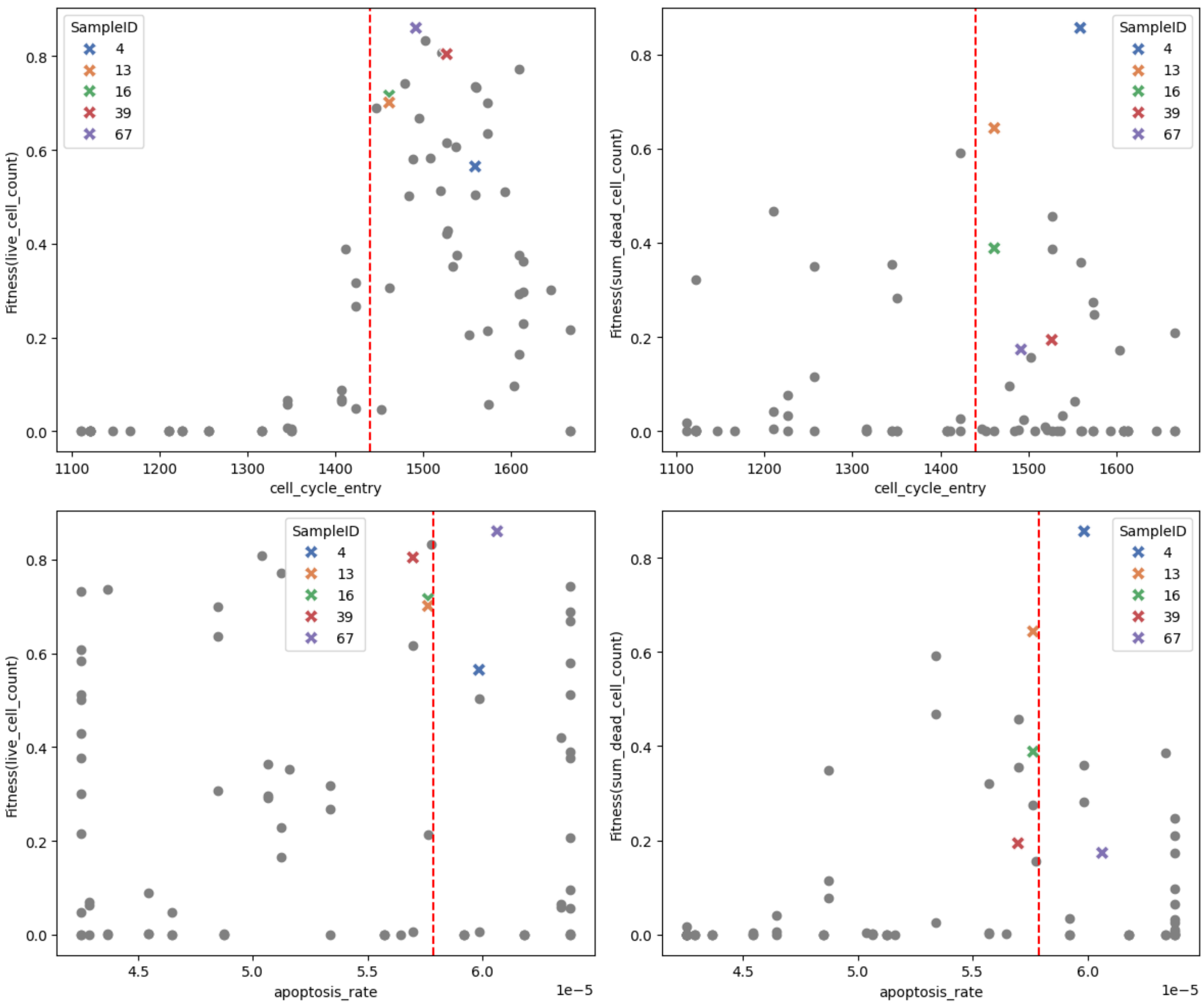


**S2 Fig. Objective function landscape and Pareto points**. Scatter plots of parameter values against the fitness scores for each Quantity of Interest (QoI). This figure illustrates the sensitivity of the fitness functions to specific parameters and identifies the Pareto points that minimize error across multiple model outputs. The dashed red lines indicate the reference values used to generate the observational data.

**S1 Table: Mechanistic rules included in the invasion model**. Each rule is defined specifically for a given cell type. Following the standardized PhysiCell grammar, each rule is modeled as a behavioral response to a microenvironmental signal. The response is mathematically defined by a Hill function, where a specific signal (e.g., local pressure or ECM density) increases or decreases a target behavior from a base value toward a saturation value. Each functional relationship is characterized by four key parameters: the base and saturation levels, the half-max, and the Hill power, which dictates the sensitivity or "switch-like" nature of the response.

| **Rule name** | **Description** |
|---|---|
| Epi. Normal CIP | Contact inhibition of proliferation in normal epithelial cells |
| EMT Normal cell | Epithelial-to-mesenchymal transition in normal epithelial cells |
| MET Normal cell | Inhibition of mesenchymal-to-epithelial transition in normal mesenchymal cells |
| Slow down N-MC | Reduced motility of normal mesenchymal cells |
| Speed up N-MC | Increased motility of normal mesenchymal cells |
| Slow down CAF | Reduced motility of cancer-associated fibroblasts |
| Speed up CAF | Increased motility of cancer-associated fibroblasts |
| Epi. Tumor CIP | Contact inhibition of proliferation in epithelial tumor cells |
| EMT Tumor cell | Epithelial-to-mesenchymal transition in tumor epithelial cells |
| MET Tumor cell | Inhibition of mesenchymal-to-epithelial transition in tumor mesenchymal cells |
| Slow down TA-MC | Reduced motility of tumor-associated mesenchymal cells |
| Speed up TA-MC | Increased motility of tumor-associated mesenchymal cells |

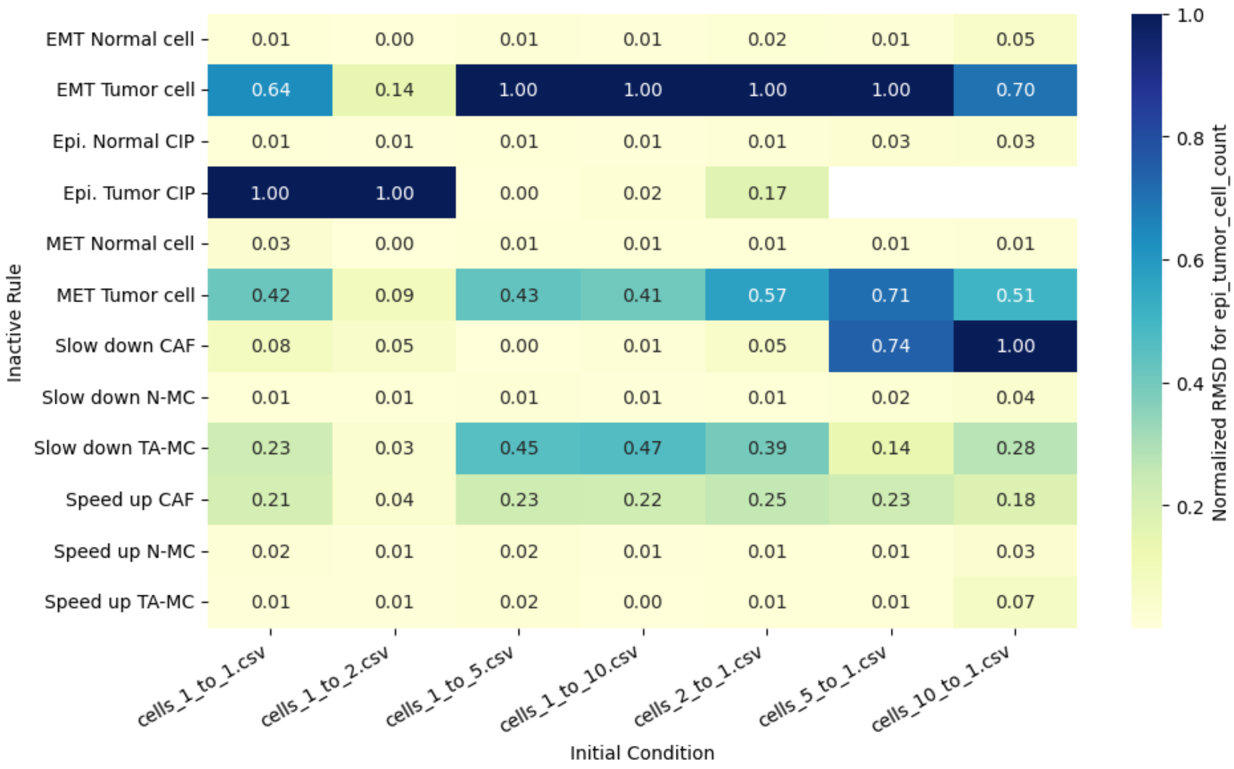


**S3 Fig. Normalized RMSD for population of epithelial tumor cells**. The normalized Root Mean Square Deviation (RMSD) between each model output under varying initial conditions and rule inactivations relative to the nominal scenario. This figure quantifies the impact of inactivating each rule on model behavior compared to the number of epithelial tumor cells in the nominal scenario.

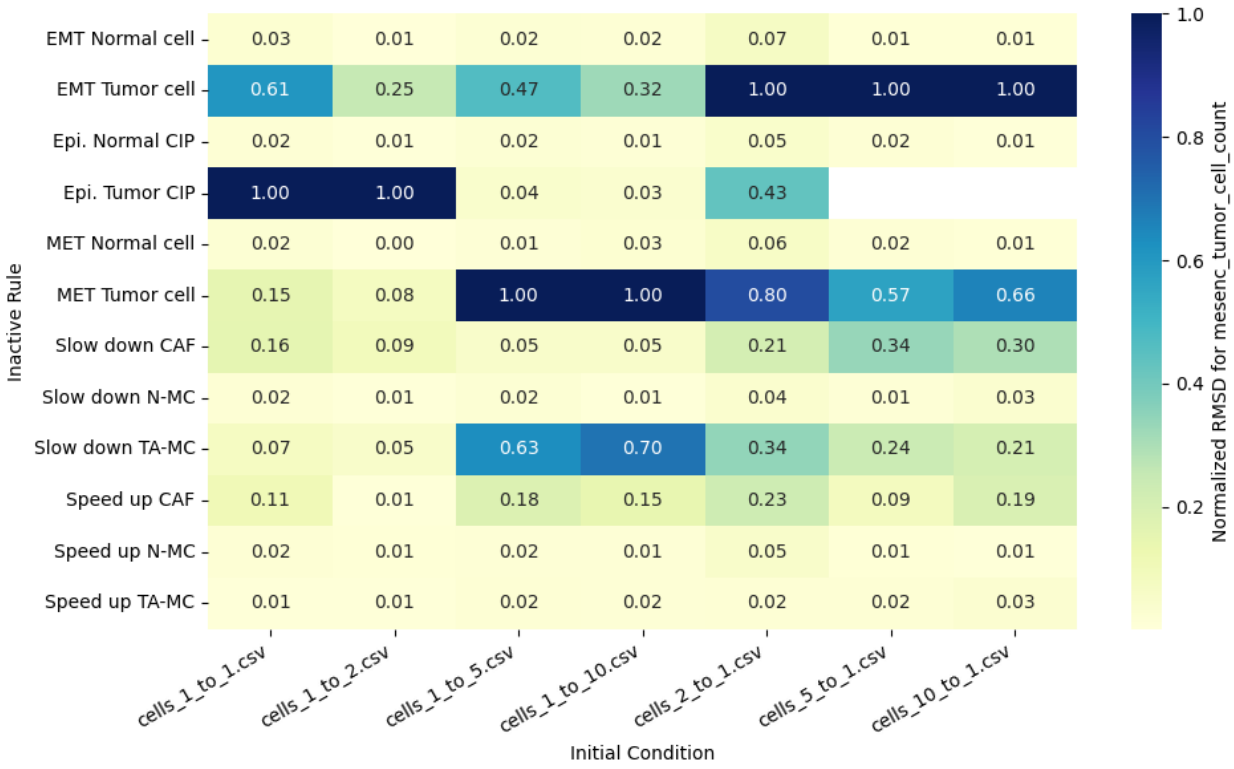


**S4 Fig. Normalized RMSD for population of mesenchymal tumor cells**. The normalized Root Mean Square Deviation (RMSD) between each model output under varying initial conditions and rule inactivations relative to the nominal scenario. This figure quantifies the impact of inactivating each rule on model behavior compared to the number of mesenchymal tumor cells in the nominal scenario.

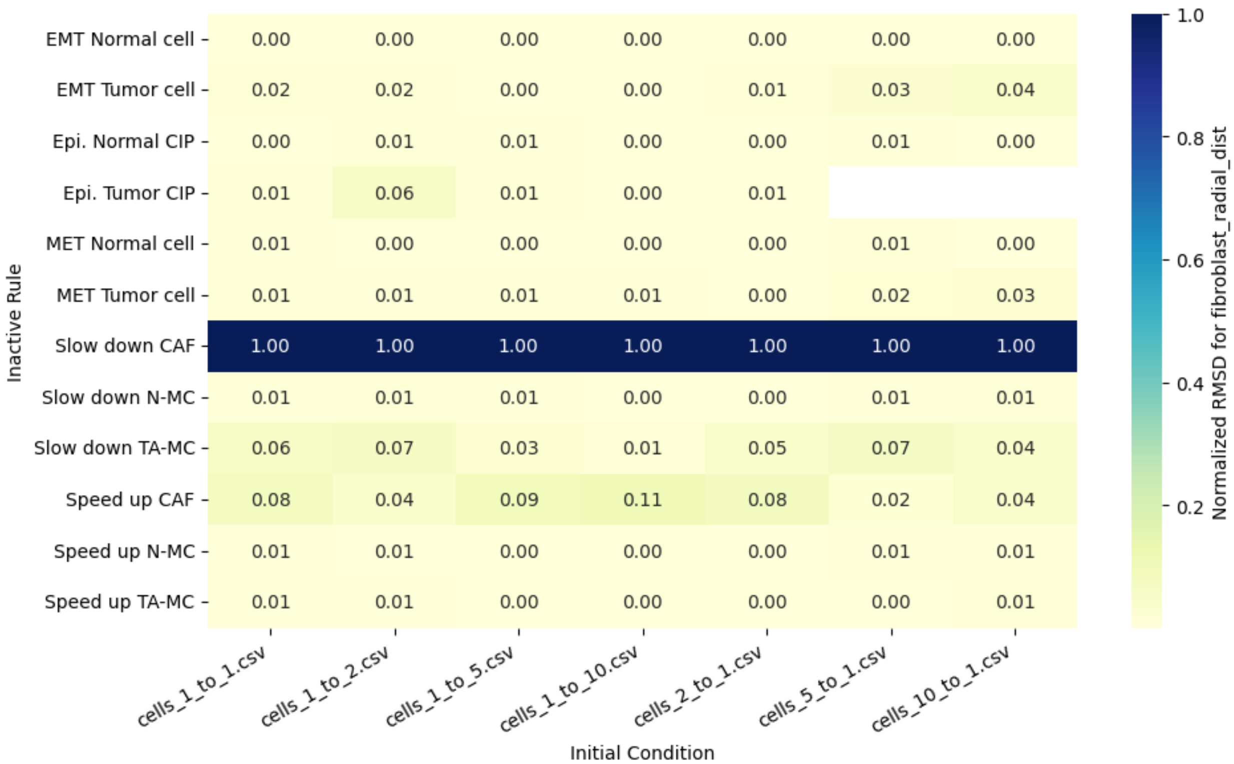


**S5 Fig. Normalized RMSD for radial distance of fibroblasts**. The normalized Root Mean Square Deviation (RMSD) between each model output under varying initial conditions and rule inactivations relative to the nominal scenario. This figure quantifies the impact of inactivating each rule on model behavior compared to the radial distance of fibroblasts in the nominal scenario.

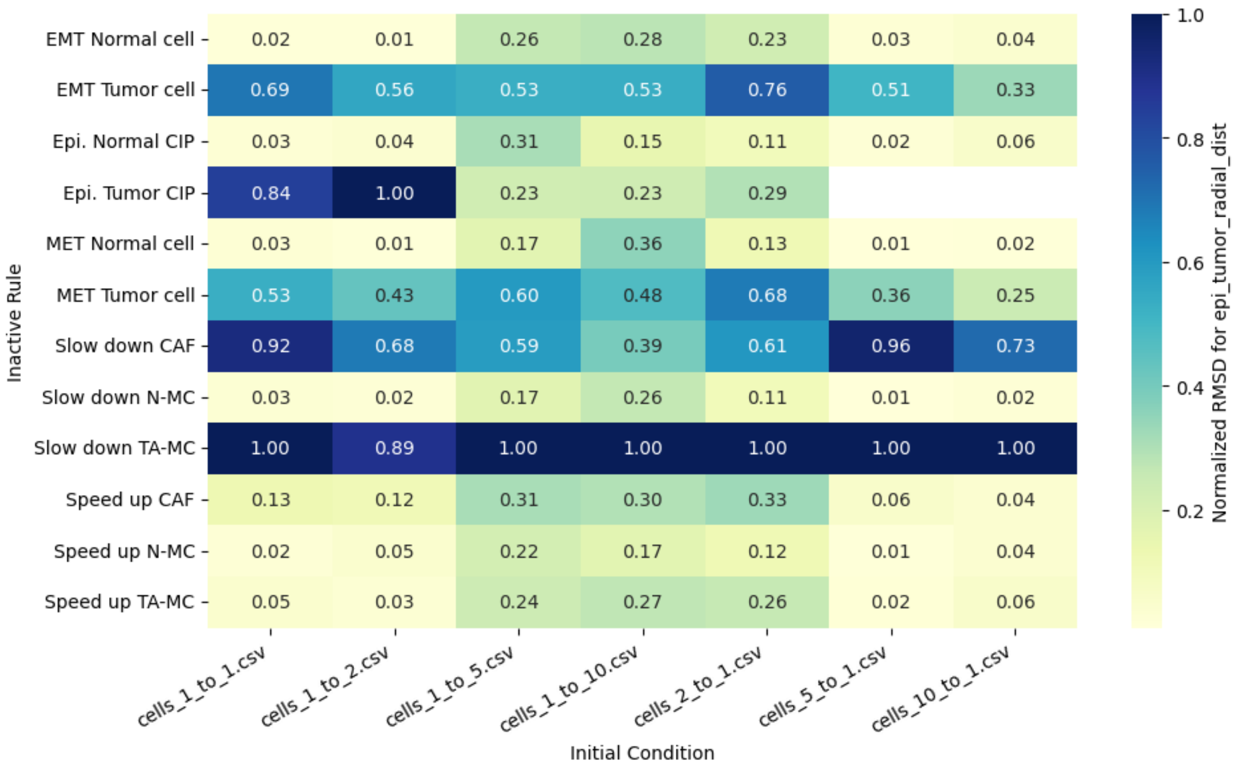


**S6 Fig. Normalized RMSD for radial distance of epithelial tumor cells**. The normalized Root Mean Square Deviation (RMSD) between each model output under varying initial conditions and rule inactivations relative to the nominal scenario. This figure quantifies the impact of inactivating each rule on model behavior compared to the radial distance of epithelial tumor cells in the nominal scenario.

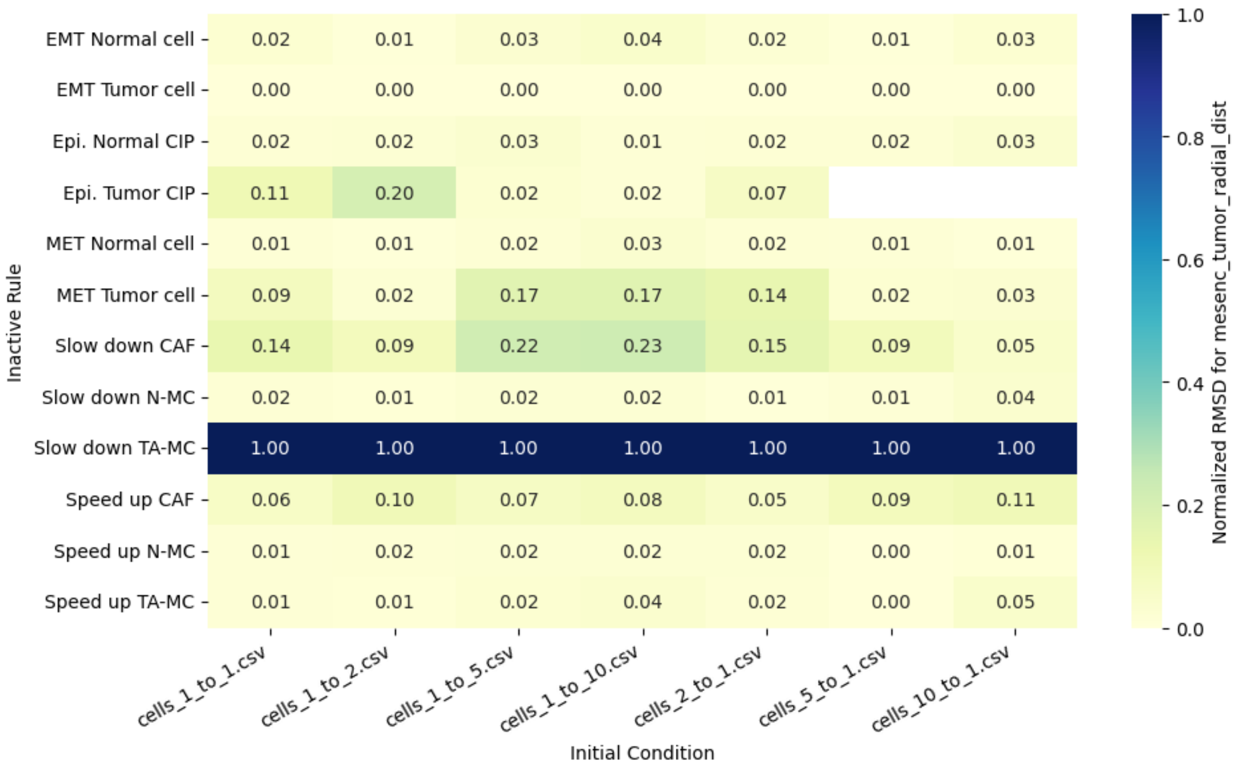


**S7 Fig. Normalized RMSD for radial distance of mesenchymal tumor cells**. The normalized Root Mean Square Deviation (RMSD) between each model output under varying initial conditions and rule inactivations relative to the nominal scenario. This figure quantifies the impact of inactivating each rule on model behavior compared to the radial distance of mesenchymal tumor cells in the nominal scenario.

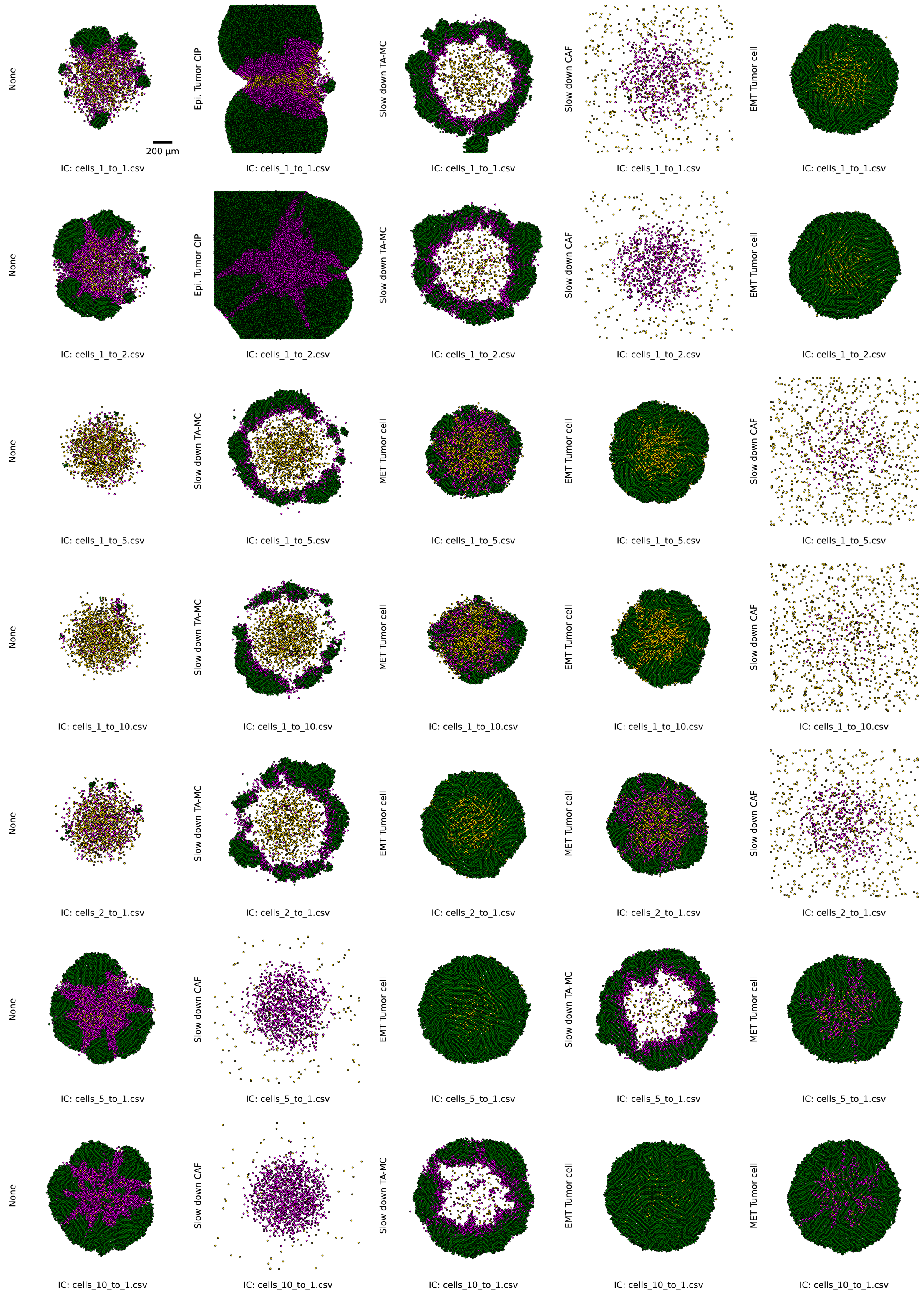


**S8 Fig. Spatial snapshots of nominal and high-impact scenarios**. Final simulation snapshots (last time point) across different initial conditions. The first column displays the nominal (baseline) behavior, while columns 2–5 visualize the scenarios with the highest impact on model outputs, ordered by their influence as determined by the sensitivity analysis.
